# Repeated multi-domain, but not single-domain, cognitive training prevents cognitive decline and amyloid pathology found in the APP^NL-G-F^ mouse model of Alzheimer disease

**DOI:** 10.1101/2020.08.24.264978

**Authors:** Jogender Mehla, Scott H. Deibel, Hadil Karem, Shakhawat Hossain, Sean G. Lacoursiere, Robert J. Sutherland, Robert J. McDonald, Majid H. Mohajerani

## Abstract

Education, occupation, and an active lifestyle, comprising enhanced social, physical, and mental components are associated with improved cognitive functions in aged people and may prevent/ or delay the progression of various neurodegenerative diseases including Alzheimer’s disease (AD). To investigate this protective effect, APP^NL-G-F/NL-G-F^ mice at 3 months of age were exposed to repeated, single- or multi-domain cognitive training. Cognitive training was given at the age of 3, 6, 9 & 12 months of age. Single-domain cognitive training was limited to a spatial navigation task. Multi-domain cognitive training consisted of a spatial navigation task, object recognition, and fear conditioning. At the age of 12 months, behavioral tests were completed for cognitive training groups and control group. After completion of behavioral testing, mice were sacrificed, and their brains were assessed for pathology. App^NL-G-F^ mice given multi-domain cognitive training compared to APP^NL-G-F^ control group showed an improvement in cognitive functions, reductions in amyloid load and microgliosis, and a preservation of cholinergic function. There were mild reductions in microglosis in the brain of APP^NL-G-F^ mice with single-domain cognitive training. These findings provide causal evidence for the potential of certain forms of cognitive training to mitigate the cognitive deficits in Alzheimer disease.

## 1. Introduction

Alzheimer’s disease (AD), the most common form of dementia, is a progressive neurodegenerative disease affecting elderly populations worldwide. AD is characterized by extracellular amyloid-beta (Aβ) plaque deposition and intracellular formation of neurofibrillary tangles (NFT), synaptic loss, and severe cognitive decline (Selkoe and Hardy, 2016; Corriveau et al., 2017). Several non-genetic risk factors such as diabetes mellitus, obesity, hypertension, brain injuries, depression or physical inactivity are related to one-third AD cases worldwide (Lye and Shores, 2000; Ownby et al., 2006; Mayeux and Stern, 2012; Norton et al., 2014). Available pharmacological therapies provide only mild and temporary symptomatic relief (Karakaya et al., 2013). Several epidemiological and clinical studies revealed that education, occupation, and physical activity can improve cognitive ability in healthy older people, and provide protection against the development and progression of AD (Friedland et al., 2001; Fratiglioni et al., 2004; Baker et al., 2010; Mayeux and Stern, 2012; Svensson et al., 2015).

Several human experimental studies have reported that cognitive and physical activities prevent AD disease progression and improve cognitive functions by reducing the cerebral Aβ plaques and amyloid angiopathy (Lazarov et al., 2005; Ambree et al., 2006; Billings et al., 2007; Mirochnic et al., 2009). Cognitive stimulation seems to improve the overall cognitive performance, especially in persons with mild-to-moderate dementia, which persisted in a 3-month follow-up. These effects are above the beneficial effects of any medication (Woods et al., 2012; Aguirre et al., 2013). Some evidence also suggests that increasing mental activity in aging people shows a favorable and novel non-pharmacological approach to prevent/ or delay age related cognitive dysfunctions and decreasing number of cases of dementia (Gates and Valenzuela, 2010). Similar effects of cognitive training have been shown in rodents. Mice exposed to social, physical, and cognitive training showed protective effect against cognitive impairment, decreased brain Aβ burden, and enhanced hippocampal synaptic immunoreactivity (Billings et al., 2007; Martinez-Coria et al., 2015). Interestingly, repeated training in the Morris water task (MWT), a spatial navigation task, also induces learning improvements for newly acquired platform locations, and reduces tau and Aβ pathology in 3xTg-AD animals (Billings et al., 2007). Martinez-Coria and colleagues also reported that recurrent training in MWT ameliorated both spatial and non-spatial forms of memory in 3xTg-AD mice (Martinez-Coria et al., 2015).

In the present study, we sought to clarify some issues concerning the effects of cognitive training on AD memory impairments and pathology. First, many of the human studies are epidemiological and, as such, are correlational in nature. Second, of the human experimental studies in which causation can be inferred many of them manipulate multiple parameters like cognitive training, physical activity, and brain pharmacology and, thus, it is difficult to ascertain which of these factors is driving the improved function and reduced pathology. A similar pattern emerges in the animal literature. Third, many of these studies investigate the effects of these lifestyle factors in aging and not AD patients specifically. These different forms of age-related cognitive decline represent different brain conditions. Fourth, studies on humans are usually confounded with a sizeable variation of socioeconomic status among participants (Yaffe et al., 2013). Fifth, the animal models of AD used in the literature vary and many of these models are not ideal for various reasons, including overexpression of protein fragments that are not found in human AD (Saito et al., 2014; Sasaguri et al., 2017). Finally, many of the studies use single-domain training paradigms, which may or not be sufficient to reduce severe memory and brain pathologies associated with AD.

In light of the aforementioned considerations, the present experiments used a new generation knock-in mouse model of AD to assess the potential positive impacts of repeated single-domain cognitive training (ST) versus multi-domain cognitive training (MT). APP^NL-G-F/NL-G-F^ were exposed to single- or multi-domain cognitive training at 3, 6, 9 & 12 months of age. Single-domain cognitive training was limited to a spatial navigation task. Multi-domain cognitive training consisted of a spatial navigation task, object recognition, and fear conditioning. At the age of 12 months, cognitive functions were assessed for all three groups. After completion of behavioral testing, mice were sacrificed, and their brains were assessed for multiple pathologies associated with AD including amyloid plaques, microglial activation, and cholinergic function. Our working hypothesis was that MT would be the most effective at preventing AD related cognitive impairments and associated brain pathologies.

## 2. Materials and Methods

### 2.1. Animals and experimental design

APP knock-in (APP-KI; App^NL-G-F/NL-G-F^) mice were provided as a gift by the laboratory of Dr. Saido at the RIKEN Center for Brain Science, Japan. A colony of these mice have been maintained at the vivarium at the Canadian Centre for Behavioral Neuroscience, University of Lethbridge. Mice were genotyped and marked using an ear notching method. In each cage, four mice were kept in a controlled environment with free access to food and water. All experimental procedures were approved by the institutional animal care committee and performed in accordance with the standards set out by the Canadian Council for Animal Care. In the present study, we used total 53 male mice (33 APP^NL-G-F^ for cognitive training; 10 APP^NL-G-F^ age match control (12 months old) & 10 C57BL/6 age match control (12 months old)) for our experiment. The mice were assigned into two experiments.

### 2.2. Experiment 1

In experiment 1 (n=16), mice were exposed to repeated multi-domain cognitive training (MT) (Figure 1). For APP-MT group, the same mice were trained on various behavioral paradigms such as the standard spatial version of the Morris water task (MWT), novel object recognition (NOR) and fear conditioning (FC) tests at 3, 6, 9 & 12 months of age. After the completion of behavioral tests, at each time point, we injected methoxy-XO4 (10 mg/kg) intraperitoneally (i.p) to 4 mice to stain amyloid plaques in their brain (Jafari et al., 2019). Then, 24 hrs after the injection, the mice were perfused and their brains were extracted for histology. We also performed immunostaining for microglial and cholinergic function.

**Fig. 1:**
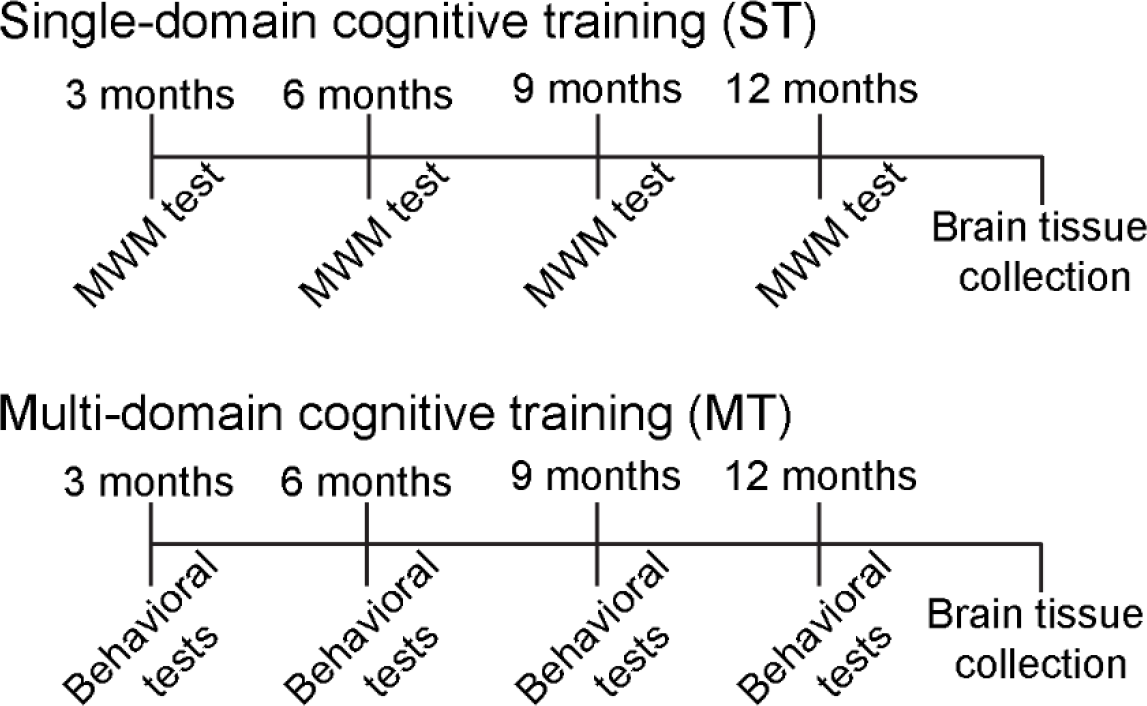
Experimental design for the study. The study was conducted for 12 months. For multidomain cognitive training (MT), we performed the spatial version of the Morris water task (MWT), novel object recognition (NOR) & fear conditioning (tone & context) tests. We used only MWT for single-domain cognitive training (ST).

### 2.3. Experiment 2

In experiment 2 (n=17), mice were given repeated single-domain cognitive training (APP-ST) using the MWT (Figure 1). This experiment was performed to rule out the effect of exercise associated with training on the MWT used in experiment 1. We hypothesized that MT used in experiment 1 would be more effective over ST used in experiment 2 to ameliorate cognitive dysfunction and AD pathology in the APP-KI mouse model. For repeated ST, the same mice were trained on the MWT at the ages of 3, 6, 9 and 12 months. After the completion of MWT probe testing at each time point, we injected methoxy-XO4 (10 mg/kg, i.p) into four mice to stain amyloid plaques in their brain (Jafari et al., 2019). Then, 24 hrs after the injection, mice were perfused after a dose of pentobarbital and their brains were extracted for histology. We also performed immunostaining for microglial and cholinergic function.

To control for the effects that repeated MT and ST every three months has on cognition and AD-related pathology, naïve groups of 12-month old APP^NL-G-F^ (APP-NT) and C57BL/6 were also included in this study.

### 2.4. Behavioral experiments

#### 2.4.1. Morris water task (MWT)

MWT experiments were conducted as reported previously (Mehla et al., 2018, 2019a,b). In brief, the pool was filled with water, made opaque by adding non-toxic white paint and water temperature was maintained at 22±1°C. The water tank was divided into four quadrants. A circular platform was kept 0.5-1 cm below the water surface in any one quadrant. Three distinct cues of different geometry were placed around the tank. On each acquisition day, mice received four training trials from each quadrant in distributed manner for eight days. The trial was completed successfully once the mouse found the platform or 60 seconds had elapsed. If the mouse failed to find the platform on a given trial, the mouse was guided to find the platform. Mice received full training again at each time-point. The latency to find the platform was used as an indicator of spatial learning ability of the mice. The swim speed was analyzed to rule out the involvement of motor function as a confounding factor. A single probe trial was conducted on ninth day to assess spatial memory performance. The data collected during the probe trial were analyzed measuring the time spent by mice in the target quadrant.

#### 2.4.2. Novel Object Recognition Test (NOR)

The test was performed as described previously (Mehla et al., 2018: 2019a). The advantage of using this test is that it does not require external motivation, reward, or punishment. It is based on the fact that mice will explore novel items. When mice are made familiar with two similar objects during the training day, they will spend more time to explore a novel object on a subsequent test day, when in a familiar environment. This pattern of behavior clearly indicates that the mice formed memories of the objects during training and noticed the presence of a novel object during testing (Antunes and Biala, 2012). A white, plastic, square box was used for the object recognition test. Briefly, mice were brought from their home cage to the experimental room and familiarized to testing box for five minutes daily for 2-3 days. Training was conducted 24 hours after the last acclimatization day. Two familiar objects were cleaned with 70% isopropyl alcohol to mask any previous order cues and allowed to dry completely. A video camera was used to record the mices’ behavior for further analysis. The mice were put into the testing box for 10 min to explore both familiar objects and recordings were made for each mouse. After 24 hrs of training, a test session was conducted. In this session, one of the familiar objects was replaced with a novel object with different geometry and texture. These objects were cleaned with 70% isopropyl alcohol and allowed to dry completely. Mice were individually placed in the testing box for 5 min to explore the objects and a recording of their behavior was made for each mouse. After each mouse completed the test, feces were removed from the testing box. The objects were also wiped with 70% isopropyl alcohol to mask the odor cues after each mouse. The data was analyzed by measuring the exploration time for familiar and novel object during training and testing days. The investigation ratio was calculated for each group.

#### 2.4.3. Fear Conditioning Test

Fear conditioning (FC) was conducted as described in previous studies (Mehla et al., 2018, 2019a). This test is conducted to assess the amygdala and hippocampus associated memory in rodents. FC was conducted in an acrylic square box. A video camera was used to record the mice’s behavior for further analysis. The floor of chamber consisted of stainless-steel rods. These rods were connected to a shock generator for the delivery of a foot-shock. A speaker was used to deliver a tone stimulus. Prior to conditioning, the chamber was cleaned with a 1% Virkon solution to mask any previous odor cues. On the conditioning day, mice were brought from their home cage into a testing room and allowed to sit undisturbed in their cages for 10 min. Mice were then kept in the conditioning square box and allowed to explore for 2 min before the onset of the tone (20 sec, 2000 Hz). In the delay conditioning test, a shock (2 sec, 0.5 mA) was given in the last two seconds of tone duration. Mice received five delayed conditioning trials, each separated by a 120-second intertrial interval (ITI). The mice were taken from the conditioning chambers one minute after the last shock and returned to their home cages. Before the tone test, the triangular chamber was cleaned with a 70% isopropyl solution. After 24 hours, the tone test was conducted in a triangular chamber that was geometrically different than the conditioning chamber to assess conditioning to the tone in the absence of the training context. For the tone test, three 20-second tones were given after a 2-minunte baseline period. Each tone presentation was separated by a 120-sec ITI. The freezing response was measured using a time sampling procedure in which an observer scored the presence or absence of the freezing response for each mouse at every two second interval. Then, for context test, mice were placed back in the original conditioning box for a 5-minute 24 hours after the tone test. During this test, freezing was scored for each mouse at every 5 second interval. Data was transformed into a percent freezing score by dividing the number of freezing observations by the total number of observations and multiplying by 100. In each conditioning and context procedures, the box was cleaned with 1% Virkon; the tone procedure box was cleaned with 70% isopropyl after each mouse trial to mask any odor cues left by the previous subject and discern context.

### 2.5. Histology

After completion of behavioral tests, mice were injected with methoxyXO4 (10 mg/kg, i.p) as described in a previous study (Jafari et al., 2019). Then, 24 hrs after injection, mice were transcardially perfused with phosphate buffered saline (PBS) followed by 4% paraformaldehyde (PFA). Brains were extracted, and post-fixed for 24 h in 4% PFA at 4°C. Later, brains were transferred to 30% sucrose after rinsed with PBS.

#### 2.5.1. Quantification of amyloid plaques in mice brain

We assessed amyloid pathology in different brain regions such as medial prefrontal cortex (mPFC), hippocampus (HPC), retrosplenial area (RSA), perirhinal cortex (PRhC) and cortical amygdalar area (CAA). Therefore, we used brain section A1 for mPFC and A2 for HPC, RSA, PRhC and CAA. Fixed brains were coronally sectioned at 40 μm using microtome. Brain sections were put on the slide and air dried for 15-20 min. Later, sections on the slides were washed twice with tri-buffered saline (TBS) for 5 min each. Then, brain sections were covered with coverslips with Vectashield H-1000 (Vector Laboratory). Finally, whole slides were imaged using NanoZoomer microscope with 20x objective magnification (NanoZoomer 2.0-RS, HAMAMTSU, JAPAN). The images were analyzed using ImageJ (US National Institutes of Health, Bethesda, Maryland) and ilastik software (Berg et al., 2019). This software automatically gives the plaques number and size corresponding with each discrete plaque. The number of plaques was quantified according to the plaque size (less than or more than 4 μm). In line with Hefendehl et al. 2011, 87% of newly generated plaques are small, having a radius of <4 μm. ImageJ (US National Institutes of Health, Bethesda, Maryland) was used to determine the plaque area.

#### 2.5.2. Immunohistochemistry for microglial and Choline acetyl transferase

Brains were serially sectioned coronally at 40□μm on a freezing microtome. Immunohistochemical procedures for IBA-1 and ChAT were performed as previously described (Saito et al., 2014; Mehla et al., 2018, 2019a). In brief, sections were fixed on positively subbed slides. Brain sections on the slides were washed in tris buffered saline (TBS), and then, blocked for 2h in TBS containing 0.3% Triton-X and 3% goat serum. The sections were incubated for 24 hours in primary antibody (prepared in TBS with 0.3% Triton-X) at room temperature in a dark humid chamber on the shaker. The following primary antibodies were used: rabbit anti-ChAT (monoclonal, Abcam, ab178850, 1:5000); Anti-Iba1 (Rabbit, SAF4318, 019-19741, Wako. Following incubation, three 10-minute washes were done, and sections were again incubated with secondary antibody for 24 hours. The following secondary antibodies were used: goat anti-rabbit-alexa-594 (IgG (H+L), A11037, Invitrogen, 1:1000 for ChAT & IBA-1). Following incubation with secondary antibody, three 10-minute washes were given. Finally, the only ChAT sections were again incubated with incubated with DAPI (1:2000 of the 20ug/ml stock in TBS) for 45-60 min. After incubation, single 5 min wash was given. Then, sections on the slides were covered with coverslips with Vectashield H-1000 (Vector Laboratory). Later, the slides were sealed with nail polish. Finally, whole slides were imaged using NanoZoomer microscope with 20x objective magnification (NanoZoomer 2.0-RS, HAMAMTSU, JAPAN). The images were analyzed using ImageJ and ilastic software (Berg et al., 2019).

### 2.6. Statistical analysis

The statistical analyses were conducted using SPSS statistical software package, version 22.0. Results are given as mean±SEM. One-way ANOVA with repeated measures was used to find statistically significant differences across the days during the acquisition phase. Paired *t*-tests were used to measure significant differences between the target and the average of other quadrants during the probe trial. Fisher’s Least Significant Difference (LSD) was used as a *post-hoc* comparison test to find significant difference between the groups. A p value < 0.05 was considered as statistically significant.

## 3. Results

### 3.1. Effect of cognitive training on various learning and memory functions of APP^NL-G-F^ mice in MWT

As can be seen in Figure 2A, in regards to 12 month old animals that were experiencing the MWT for the first time, the C57BL/6 (control) group learned to find the hidden platform location with training while the APP^NL-G-F^ (APP-NT) group did not. Consistent with these observations, control mice (12 months old) showed significant (p<0.01; n=10 mice) decreases in escape latency on day 8 (13.23±1.91 sec) as compared with day 1 (39.60±2.37 sec), indicating that these mice learned to swim to the goal efficiently (Figure 2B & Supplementary Fig 1A). The APP-NT group exhibited impairments in learning the spatial task as revealed by a nonsignificant difference in the escape latency on day 8 (33.31±3.25 sec) in comparison to day 1 (39.35±3.94 sec; n=10 mice Figure 2A-B & Supplementary Fig 1A).

**Fig. 2:**
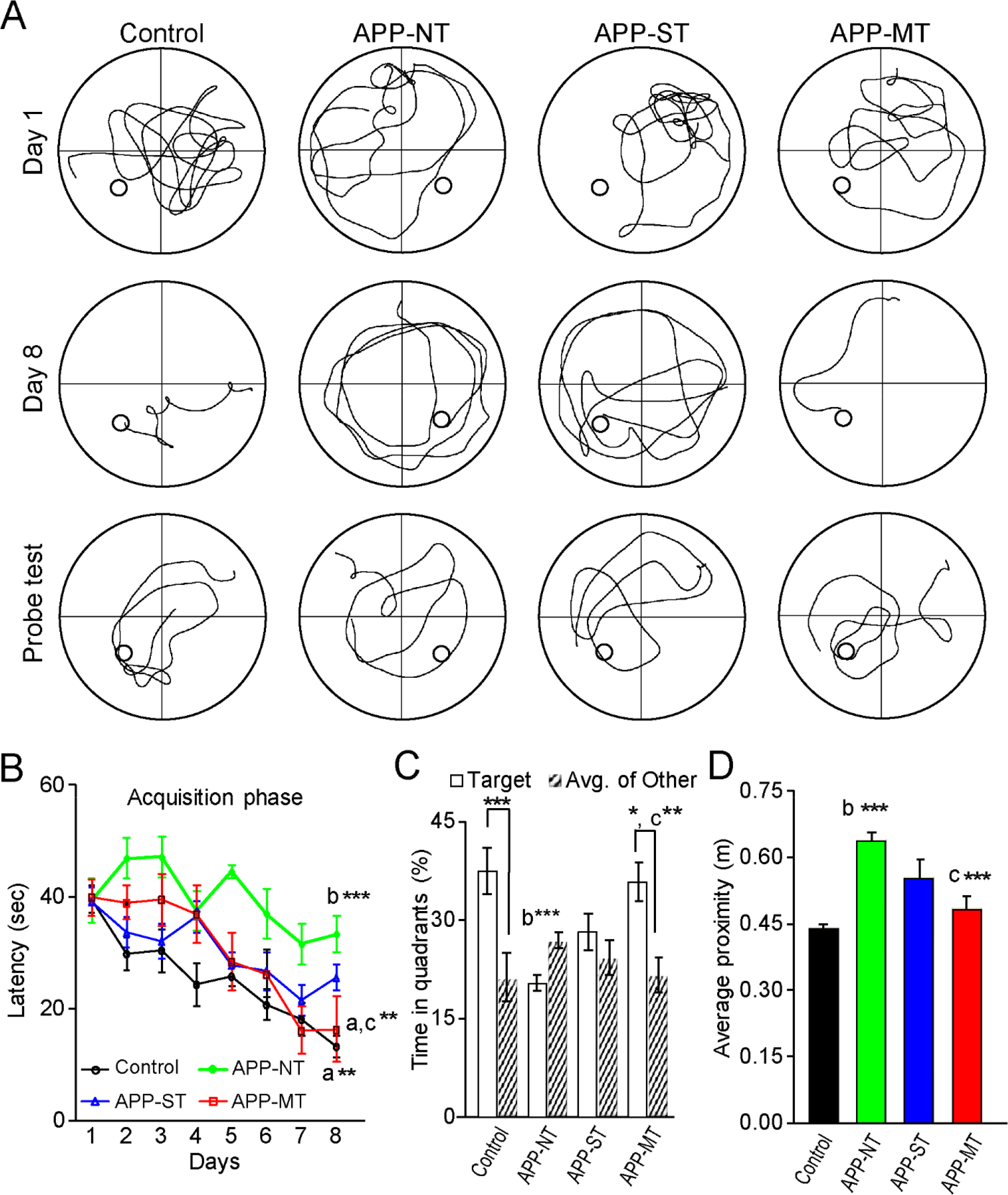
Effects of cognitive training on spatial learning and memory functions of 12 months old control and App^NL-G-F^ mice in the MWT. (A) Representative swim path on Day 1, Day 8 and probe trial for various experimental groups. Overall comparison of experimental groups considering mean latency to find the hidden escape platform during the acquisition phase. (B) Mean latency to find the hidden escape platform during the acquisition phase for various groups. (C) Percent time spent by mice in target quadrant and average of other quadrants during the probe trial. (D) Average proximity to platform during probe trial. Data is presented as mean ± SEM. *p < 0.05; **p < 0.01; ***p < 0.001; a- as compared with day 1 for group control, APP-MT, b- as compared with control group on day 8; c- as compared with APP-NT group; Control group- C57BL/6. APP-NT APP^NL-G-F^ mice with no training. APP-ST group- APP^NL-G-F^ mice exposed to single-domain cognitive training (ST). APP-MT group- APP^NL-G-F^ mice exposed to multi-domain cognitive training (MT).

The single domain cognitive training group (APP-ST) was impaired in spatial learning compared to the multiple domain cognitive training (APP-MT) group (Figures 2AB & 3B). No significant difference was observed in escape latency for the APP-ST group between day 1 (39.05±3.25 sec) and 8 (25.52±5.84 sec; n=8 mice) showing no improvement in learning ability of these mice given this nonpharmacological treatment condition (Figure 2B & Suppl. Figure 1B). Whereas, the APP-MT group, however showed spatial learning abilities similar to the control group. Consistent with this claim, there was a significant decrease in the escape latency on day 8 (16.29±2.02 sec) when compared with day 1 (39.88±3.46 sec; p<0.01; n=7 mice; Figures 2B & Suppl. Figure 1C) for the APP-MT animals. A one way ANOVA revealed a significant (*F*(_3,31_)=7.297, p=0.001) difference among the experimental groups. In comparison to the control group, the APP-NT group took more time to reach the platform on day 8 which is indicated by significant (p<0.001) increased escape latency (13.23±1.91 sec for control group vs. 33.31±3.25 sec for the APP-NT group; Figure 2B). Mice exposed to ST did not show significant difference in the escape latency on day 8 when compared with the APP-NT group (Figures 2B & Suppl. Figure 1B) showing no improvement in learning function of mice due to single-domain training. However, the APP-MT group showed significant (p<0.01) decrease in escape latency on day 8 in comparison to the APP-NT group indicating an improvement in learning performance of mice due to cognitive training (Figures 2B & Suppl. Figure 1C). We did not find any significant differences in the swim speed of experimental groups at testing age points (Suppl. Figure 2).

**Fig. 3:**
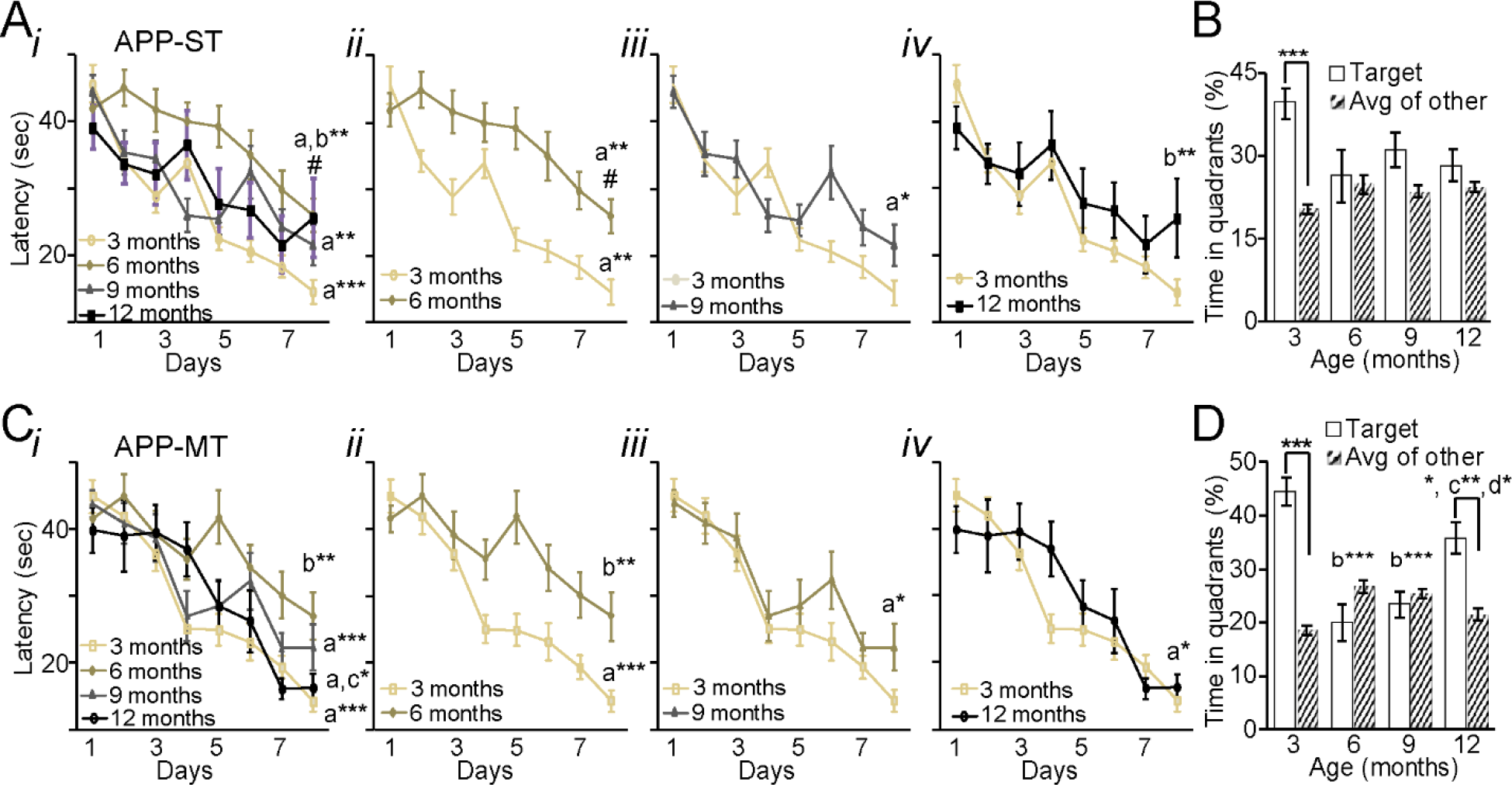
Effect of repeated single and multi-domain cognitive training (ST, and MT) on spatial learning and memory functions of 3, 6, 9, &12 months old APP^NL-G-F^ mice as assessed with the spatial version of the MWT. (A) i. Mean latency to find the hidden escape platform during the acquisition phase examined at 3, 6, 9, &12 months old ÄPP^NL-G-F^. ii. Comparison of escape latency of 3- and 6-months old APP mice exposed to ST. iii. Comparison of escape latency of 3- and 9-months old APP mice exposed to ST. iv. Comparison of escape latency of 3- and 12-months old APP mice exposed to ST. (B) Percent time spent by mice in target quadrant and average of other quadrants during the probe trial. a- as compared with day 1 for 3, 6, & 9 months old mice; b- as compared with 3 months old mice on day 8; #- as compared to 3 months old mice, ***- as compared to 3 months old. APP-ST group- APP^NL-G-F^ mice exposed to single-domain cognitive training (ST). (C-D) Same as A and B for APP mice exposed to MT. a- as compared with day 1 for 3, 9, & 12 months old mice; b- as compared with 3 months old mice; c- as compared with 6 months old mice; d- as compared to 9 months old. APP-MT group- App^NL-G-F^ mice exposed to multi-domain cognitive training (MT). Data is presented as mean±SEM. **p < 0.01; ***p < 0.001.

24 hrs after the last trial on day 8 of the acquisition phase, a probe trial was conducted to assess the spatial memory of the 12-month mice assigned to the different experimental groups. We found that mice in control group spent significantly (p<0.001; n=10 mice) more time in the target quadrant in comparison to average of other quadrants (Figure 2C), indicating normal retention of the spatial memory acquired during MWT training. However, no significant difference was found in the time spent in the target versus the average of the other quadrants by mice in the APP-NT group indicating an impairment in spatial memory retention (Figure 2C). Similarly, the APP-ST group did not show good spatial memory retention as indicated by nonsignificant difference among the time spent in the target versus the average of the other quadrants (Figure 2C). In contrast, the MT condition ameliorated the memory deficit in APP^NL-G-F^ mice, indicated by the fact that the time spent in target quadrant by mice was significantly (p<0.05; n=7 mice) more than the time spent an average of the other quadrants (Figure 2C). One way ANOVA showed a significant (*F*(_3,31_)=8.995, p=0.005) difference in the time spent in the target quadrant amongst the experimental groups. LSD *post-hoc* analysis revealed that the APP-NT group spent significantly less time (p<0.001; 37.41±3.49% for control group vs. 20.24±1.17% for the APP-NT group) in the target quadrant when compared with control group (Figure 2C). Significant increases in time spent in the target quadrant by the APP-MT group was found when compared with the mice in APP-NT group (Figure 2C) and no significant difference was found between time spent in target quadrant by mice in ST and APP-NT groups (Figure 2C). Similarly, average proximity was significantly (p<0.001) higher in APP-NT group as compared to control group indicating diffuse pattern of target search during probe trial (Figure 2D). MT (p<0.01) group spent mostly time near to platform zone as evidenced by significantly less average proximity in comparison with APP-NT group alone whereas no significant difference was found between APP-NT and ST groups (Figure 2D).

We also compared the data from the repeated ST and MT groups in an age dependent manner. In the ST group, 3 months old mice took significantly less time to find the platform on day 8 as compared to day 1 (p<0.001, Figure 3Ai-ii). Six and nine months old APP^NL-G-F^ mice in ST group also learned the task to some extent as indicated by significantly lower latency on day 8 when compared to day 1 (Figure 3Ai-iii), however, 6 months old APP^NL-G-F^ mice took significantly more time to reach the platform on day 8 in comparison to 3 months old (p<0.01). This pattern of results indicates impaired spatial learning ability of 6 months old APP^NL-G-F^ mice in APP-ST group. Furthermore, 12 months old APP^NL-G-F^ mice in APP-ST group did not appear to learn the task as no significant difference was found in the escape latency on day 8 and day 1 (Figure 3Ai,iv). Moreover, these mice also took significantly more time to reach the platform on day 8 when compared with 3 months old (p<0.01). We did not find any significant difference in the swim speed of experimental mice throughout the different testing age points (Suppl. Figure 3). In the APP-ST group, only the 3 months old mice spent significantly more time in the target quadrant compared with the average of the other quadrants indicating normal spatial memory function (p<0.001; n=17 mice; Figure 3B). However, no significant difference was found in the time spent in target quadrant and average of other quadrants by 6, 9- and 12-months old mice showing impaired spatial memory (Figure 3B) in these age groups.

Three months old APP^NL-G-F^ mice with MT learned the MWT effectively as indicated by significantly (p<0.001; n=16 mice) decreased in escape latency on day 8 when compared to day 1 (Figure 3Ci-ii). However, 6 months old mice in same group did not learn the task as indicated by non-significant difference in the latency on day 1 and day 8 (n=12 mice). Additionally, these 6 months old mice also took significantly (p<0.01) more time to reach the platform on day 8 in comparison with 3 months old mice (Figure 3Ci-ii). We found some improvement in learning behavior of 9 months old APP mice exposed to MT as these mice took significantly less time to find the platform on day 8 when compared to day 1 (p<0.05; n=9 mice; Figures 2Ci,iii). Furthermore, 12 months old APP^NL-G-F^ mice with MT showed a clear improvement in learning ability evidenced by significantly (p<0.05; n=7 mice) lower escape latencies on day 8 in comparison to day 1 (Figures 3Ci,iv). We did not find any significant difference in the swim speed of experimental mice throughout the different testing age points (Suppl. Figure 3). For the probe trial, a one-way ANOVA revealed a significant (*F*(_3,42_)=16.108, p < 0.001) difference between different age testing points for the APP-MT group. LSD *post hoc* analysis indicated that 6 and 9-month-old mice in the APP-MT group spent significantly less time in the target quadrant when compared to 3 months old mice showing that the older mice from this group are showing an age-related deficit in spatial memory retention (Figure 3D). However, 12 months old mice with MT showed an improvement in memory retention evidenced by significantly more time spent by 12 months old mice in comparison to the 6 (p<0.01) and 9 (p<0.05) months old groups (Figure 3D). Taken together, these results clearly show that MT protects the learning and memory network centered on the hippocampus from AD-related brain changes which normally lead to impairments in the spatial version of the Morris water task (Mehla et al., 2019a).

### 3.2. Effect of cognitive training on the memory function of App^NL-G-F^ mice in a novel object recognition test

As can be seen in Figure 4B, control mice showed normal novel object recognition (NOR) memory whereas APP-NT mice (12 months) did not. During the training, no significant difference was found in exploration time for object 1 and 2 in any experimental groups (Suppl. Figure 4). However, control mice explored a novel object for significantly (p<0.01) longer time when compared to the familiar object (Suppl. Figure 4B). Furthermore, control mice showed a significantly (p<0.01; n=10 mice) higher investigation ratio for the novel object in comparison to the familiar object indicating these mice are spending more time exploring the novel object while the APP-NT group did not, suggesting that the latter group had an object memory impairment (Figure 4B). APP^NL-G-F^ mice (12 months old) with ST did not show an improvement in memory function evidenced by no significant difference in the investigation ratio for novel and familiar object (Figure 4B). In contrast, APP^NL-G-F^ mice (12 months) exposed to MT showed significantly (p<0.001; n=7 mice) higher investigation ratio for novel object when compared with familiar object which is also evidenced by significantly more exploration time for a novel object than familiar (Suppl. Figure 4B). We also found that 6 and 9 months old APP^NL-G-F^ mice exposed to repeated ST or MT did not show an improvement in object memory function in this task (Data not shown). Since this task has been previously shown to be dependent on the perirhinal cortex (Antunes and Biala, 2012; Watson and Lee, 2013) it appears that MT can reverse dysfunction of this medial temporal region following repeated training episodes during aging.

**Fig. 4:**
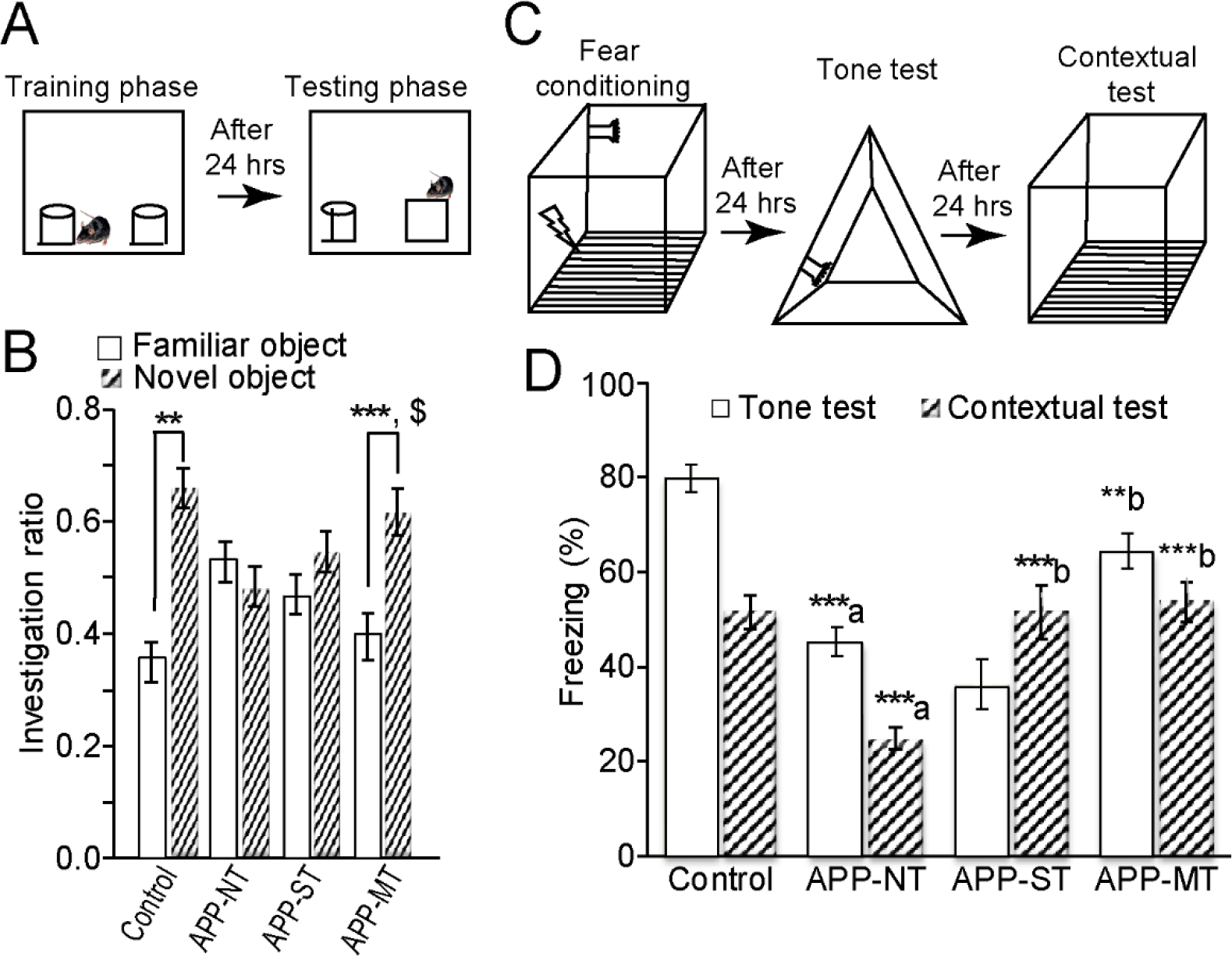
Effect of cognitive training on learning and memory functions of 12 months old APP^NL-G-F^ mice in the novel object recognition (NOR) and fear conditioning tests. (A) Schematic representation of the novel object recognition test. (B) Investigation ratio for familiar and novel object. (C) Schematic representation of the fear conditioning test. (D) Percent freezing for tone and contextual test. Data is presented as mean ± SEM. *p < 0.05; **p < 0.01; ***p < 0.001; a- as compared with control group. b- as compared with APP-NT group. Control group- C57BL/6. APP-NT group- App^NL-G-F^ with no training. APP-MT group- App^NL-G-F^ mice exposed to multidomain cognitive training (MT). APP-ST group- App^NL-G-F^ mice exposed to single-domain cognitive training (ST).

### 3.3. Effect of cognitive training on memory function of APP^NL-G-F^ mice in fear conditioning

Figure 4D shows the results of the fear conditioning to a discrete auditory cue. Control mice showed conditioned fear to an auditory cue and context associated with a fearful stimulus (foot-shock) while the APP-NT mice (12 months old) did not. Consistent with these observations, APP-NT mice (12 months old) showed significantly (p<0.001) less percentage of freezing to the tone (79.67±2.83% and 45.33±2.95% for control and APP-NT groups, respectively) and context (51.50±3.46% and 24.83±2.39% for control and APP-NT groups, respectively) tests when compared with the control mice (Figure 4D). APP^NL-G-F^ mice (12 months old) exposed to MT showed an improvement in memory function indicated by significantly increases in percentage of freezing to a conditioned tone (p<0.01) and context (p<0.001) in comparison with the APP-NT group (Figure 4D). The group receiving ST also showed an increase in percentage of freezing of APP-NT mice (12 months old) to the conditioned context, however, no improvements in fear conditioning to the tone was found (Figure 4D). We also found that 6 and 9 months old APP^NL-G-F^ mice exposed to MT or ST did not show any improved performance on either component of fear conditioning (Data not shown).

This pattern of data suggests that the neural networks mediating fear conditioning are compromised in the APP^NL-G-F^ mouse model of AD and that repeated MT protects these brain areas from associated AD pathology.

### 3.4. Effect of cognitive training on amyloid pathology of APP^NL-G-F^ mice

We assessed amyloid pathology in different brain regions such as mPFC, HPC, RSA, PRhC and CAA. Therefore, we used brain section A1 (bregma +1.94) for mPFC and A2 (bregma -3.08 mm) for HPC, RSA, PRhC and CAA. Figures 5B show a representative distribution of Aβ plaques size by total number of plaques in brain sections of various experimental groups. Lowest distribution of Aβ plaques size by total number of plaques was observed in APP^NL-G-F^ mice exposed to MT. Additionally, MT significantly decreased amyloid load in the brain of APP^NL-G-F^ mice, however, ST did not (Figure 5A-B). We observed significant decreases in percent area stained with Aβ plaques in both brain sections of mice in the MT group in comparison to the APP-NT group (Figure 5C). LSD posthoc analysis also indicated that APP^NL-G-F^ mice exposed to repeat MT showed significantly [p<0.05 for sections A1 (bregma +1.94 mm) & A2 (bregma -3.08 mm)] less Aβ plaques burden in both A1 and A2 brain sections when compared to APP-NT group (Figure 5 C). Though, APP^NL-G-F^ mice in the ST group also showed decrease in Aβ plaques deposition in both brain sections, the differences were not statistically significant (Figure 5B-C).

**Fig. 5:**
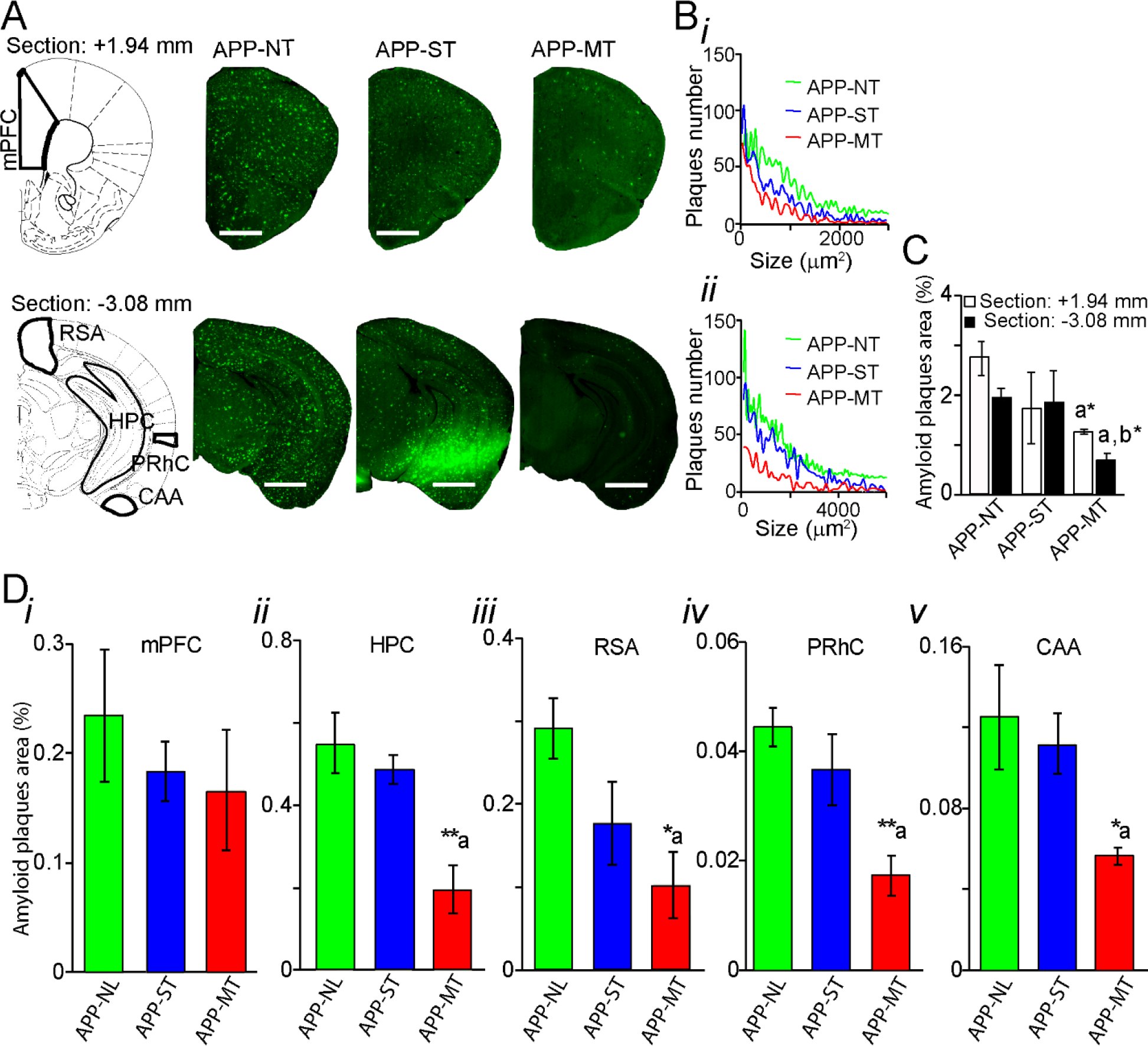
Amyloid plaque distribution in brain of 12 months old App^NL-G-F^ mice. (A) Photomicrographs of amyloid plaques stained with methoxy-XO_4_ (green) in two brain sections (top: bregma +1.94 mm; bottom: -3.08 mm). (B) i-ii Corresponding distributions of plaque size by total number of plaques for each brain section shown in A. (C) Summary results of total amyloid plaques number in brain. (D) Amyloid plaque area in medial prefrontal cortex (mPFC)(i), hippocampus (HPC)(ii) retrospenial area (RSA)(iii), perirhinal cortex (PRhC)(iv), and cortical amygdalar area (CAA)(v). Scale bars represent 1 mm for section +1.94 mm and 2.5 mm for section -3.08 mm. Data is presented as mean ± SEM. *p < 0.05, **p < 0.01; a- as compared to APP-NT group. b- as compared to APP-ST group. APP-NT group- App^NL-G-F^ with no training. APP-ST group- App^NL-G-F^ mice exposed to single-domain cognitive training (ST). APP-MT group- APP^NL-G-F^ mice exposed to multi-domain cognitive training (MT).

We also investigated the deposition of Aβ plaques in various brain regions such as mPFC, HPC, RSCA, PRhC and CAA as these brain regions are involved in various cognitive functions including learning and memory (Figures 5D). App^NL-G-F^ mice with MT showed a significant decrease in Aβ plaques burden in HPC (p<0.01, Figure 5Dii), RSA (p<0.05, Figure 5Diii), PRhC (p<0.01, Figure 5Div) and CAA (p<0.05, Figure 5Dv) when compared to mice in the APP-NT group. However, ST did not cause significant change in the Aβ deposition in these brain regions of App^NL-G-F^ mice when compared to APP-NT group (Figure 5D). None of these interventions significantly decreased plaques burden in mPFC when compared with the APP-NT group (Figure 5Di). We did not perform Aβ staining in 6 and 9 months old App^NL-G-F^ mice with MT and ST as no improvement in cognitive functions was found at this age.

Additionally, we also assessed amyloid pathology in medial septum-diagonal band complex (MSDB), a prominent region for cholinergic inputs to various brain regions especially HPC. Figure 7A-B exhibit a representative distribution of Aβ plaques size by total number of plaques in MSDB complex of various animal groups. App^NL-G-F^ mice exposed to MT showed lowest distribution of Aβ plaques size by total number of plaques (Figure 7B). APP-MT & APP-ST groups showed significant (p<0.01 for ST & p<0.001 for MT, respectively) reduction in Aβ plaques number, however, percent Aβ plaques area was significantly decreased in the MT (p<0.01) group when compared to APP-NT group alone (Figure 7D).

The results of repeated cognitive training on amyloid pathology in the App^NL-G-F^ mouse was clear. Amyloid pathology was drastically reduced in brain areas implicated in the learning and memory functions assayed in these same subjects. Namely, amyloid pathology was reduced in the HPC, RSC, PRhC, CAA, and the MSDB regions.

### 3.5. Effect of cognitive training on brain microgliosis of APP^NL-G-F^ mice

Figure 6 indicates increased microgliosis in the brain of APP-NT mice when compared with control group. Overall, MT decreased microgliosis in the brains of the App^NL-G-F^ mice exposed to this treatment in comparison with App^NL-G-F^ mice that were not given cognitive training (Figure 6A). We also found significant (p<0.001) increase in activated IBA-1, a marker of microgliosis in brain sections A1 & A2 of the APP-NT group as compared to control (Figure 6A-B). However, ST reduced percent microgliosis only in brain section A1 (Figure 6B). APP^NL^ ^G-F^ mice that experienced MT showed significant decrease in microgliosis in comparison with mice in APP-NT group (Figure 6B).

**Fig. 6:**
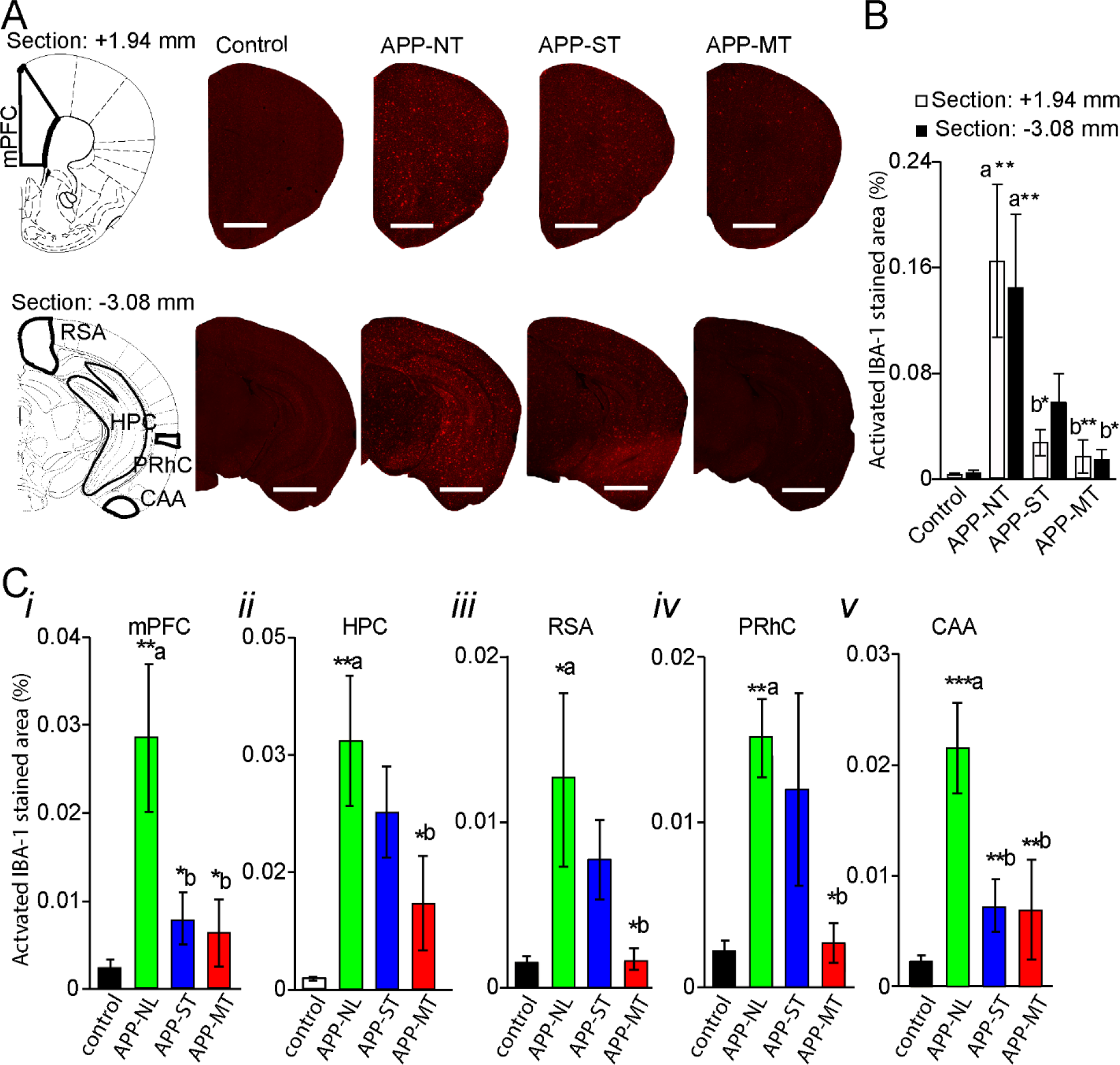
Microgliosis in brain of 12 months old mice. (A) Photomicrographs of activated IBA-1, a marker of microgliosis in brain. Representative of activated microglia in two brain sections (top: bregma +1.94 mm; bottom: -3.08 mm). Scale bars represent 1 mm for section +1.94 mm and 2.5 mm for section -3.08 mm. (B) Activated IBA-1 immunostained area in brain sections +1.94 mm and section -3.08 mm. (C) Microgliosis in different brain regions of 12 months old mice. Activated IBA-1 immunostained area in medial prefrontal cortex (mPFC)(i), hippocampus (HPC)(ii), retrospenial area (RSA)(iii), perirhinal cortex (PRhC)(iv), and cortical amygdalar area (CAA). Data is presented as mean ± SEM. *p < 0.05, **p < 0.01, ***p<0.001, a- as compared to control group; b- as compared to APP-NT group. Control group- C57BL/6. APP-NT group- App^NL-G-F^ with no training. APP-MT group- App^NL-G-F^ mice exposed to multi-domain cognitive training (MT). APP-ST group- App^NL-G-F^ mice exposed to single-domain cognitive training (ST).

We also quantified the microgliosis in different brain regions which are involved in various cognitive functions. Percent area immunostained by IBA-1 was significantly higher in the APP-NT group in comparison with control mice (Figures 6C). Furthermore, we also observed significantly increases in microgliosis in mPFC (p<0.01, Figure 6Ci), HPC (p<0.01, Figure 6Cii), RSA (p<0.05, Figure 6Ciii), PRhC (p<0.01, Figure 6Civ) and CAA (p<0.001, Figure 6Cv) brain regions of the APP-NT group in comparison with controls. A significant (p<0.05) reduction in microgliosis was also observed in mPFC (p<0.05, Figure 6Ci), and CAA (p<0.01, Figure 6Cv) brain regions of the ST group when compared with the APP-NT group. The MT group showed statistically significant reductions of microgliosis in mPFC (p<0.05), HPC (p<0.05), RSC (p<0.05), PRhC (p<0.05) and CAA (p<0.01) brain regions when compared to the APP-NT group (Figures 6Ci-v).

We also assessed microgliosis in MSDB complex. The APP-NT group showed increases in microgliosis in MSDB complex evidenced by increased activation of IBA-1 and percent area stained by IBA-1 when compared to control group (Figures 7E,F,G). MT groups showed significant reduction in microgliosis (p<0.01) in MSDB complex in comparison with APP-NT group, however, ST intervention did not cause any reductions in microgliosis in this brain region (Figures 7 F,G).

**Fig. 7:**
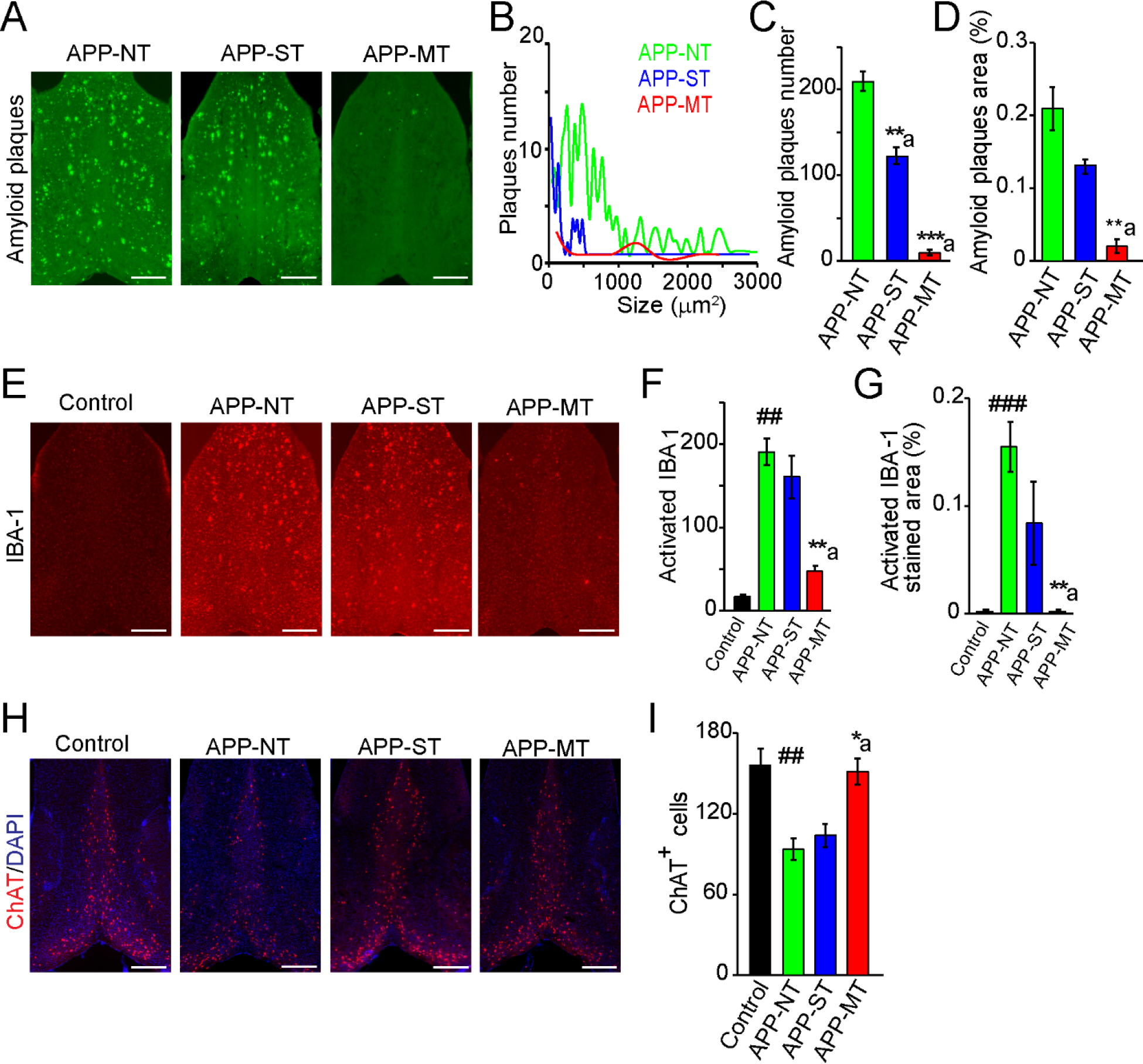
Amyloid pathology, microgliosis and cholinergic function in the medial septum-diagonal band (MSDB) complex of the basal forebrain of 12 months old mice. (A) Photomicrographs of amyloid plaques stained with methoxy-XO_4_ (green) in MSDB complex. (B) Corresponding distributions of plaque size by total number of plaques. (C) Total amyloid plaques number in MSDB complex. (D) Amyloid plaque area in MSDB complex. (E) Photomicrographs of immunostaining of activated IBA-1 (red) in MSDB complex. (F) Activated IBA-1 number in MSDB complex. (G) Activated IBA-1 immunostained area in MSDB complex. (H) Photomicrographs of immunohistochemistry staining of choline acetyl transferase (ChAT) in MSDB complex. ChAT- a cholinergic marker stained with monoclonal rabbit anti-ChAT antibody (red), DAPI-stained the nuclei (blue). (I) Quantification of ChAT in MSDB complex. Scale bar- 500 μm for whole sections. Data is presented as mean±SEM. *p < 0.05, **,##p < 0.01, ***,###p < 0.001, a- as compared to APP-NT group; #- as compared to control group. Control group- C57BL/6. APP-NT group- App^NL-G-F^ with no training. APP-MT group- APP^NL-G-F^ mice exposed to multi-domain cognitive training (MT). APP-ST group- App^NL-G-F^ mice exposed to single-domain cognitive training (ST).

In summary, APP^NL-G-F^ mice showed elevated microgliosis in the same brain regions that are functionally compromised in this AD mouse model as revealed via behavioural analysis. Further, the results showed that MT reduces these elevated microgliosis activation levels in these brain regions. However, ST reduces microgliosis only in mPFC and CAA.

### 3.6. Effect of cognitive training on cholinergic function of APP^NL-G-F^ mice

To assess cholinergic function, we performed brain immunostaining for ChAT. Mice in the APP-NT group showed decrease in ChAT positive cells in MSDB complex when compared with control mice (Figure 7H,I). Figure 7I showed that the MT intervention prevented the reduction in ChAT positive cells in MSDB complex in comparison with APP-NT mice. Consistent with these observations, one way ANOVA revealed significant (*F*(_3,11_)= 8.929, p= 0.003) difference amongst the experimental groups. LSD posthoc analysis revealed a significant (p<0.01) decrease in ChAT positive cell in MSDB complex of the APP-NT group (93.73±12.60) when compared with the control group (156.12±7.88, Figure 9I). The APP-MT group showed a reduction in the loss of cholinergic cells (p<0.05) in the MSDB complex in comparison to the APP-NT group (Figure 7I). Conversely, the APP-ST group did not show significant (p>0.05) difference in number of ChAT positive cells when compared with the APP-NT group (Figure 7I). We did not perform ChAT immunostaining in 6 and 9 months old App^NL-G-F^ mice with MT or ST as these mice did not show improvement in cognitive functions at this age.

Consistent with our previous work (Mehla et al., 2019a), App^NL-G-F^ mouse model of AD shows drastic reductions in cholinergic neurons in the MSDB. Importantly, the effects of cognitive training on the ascending cholinergic system that innervates the HPC were clear. MT but not ST prevented cholinergic cell loss in the MSDB complex. The resulting enhanced cholinergic tone in HPC might be responsible for at least some of the functional recovery found following MT in this AD mouse model like improvements in spatial learning and memory functions.

## 4. Discussion

In this series of experiments, we sought to assess the potential of cognitive training as a preventative treatment for cognitive impairments and associated brain pathology in AD. Specifically, we tested the effects of repeated exposure to different types of cognitive training in a new generation of mouse model of AD (APP^NL-G-F^). Male APP^NL-G-F^ mice at 3 months of age were exposed to single-domain or multi-domain cognitive training. ST consisted of a water-based spatial navigation task. MT consisted of the same spatial navigation task as well as novel object recognition and fear conditioning. These different forms of cognitive training were given repeatedly at the age of 3, 6, 9 & 12 months of age. At the age of 12 months mice were sacrificed and their brains assessed. APP^NL-G-F^ mice given MT compared to ST showed dramatic improvements in cognitive functions. MT also reduced brain pathology associated with AD including amyloid load and microgliosis as well as preservation of cholinergic function in the brains of App^NL-G-F^ mice. Taken together, this pattern of results provides important experimental evidence that repeated MT can reverse cognitive deficits and associated brain pathology found in Alzheimer disease.

### 4.1. Strengths of the current approach

The present series of experiments is unique in several ways that may represent an advancement in our understanding of the beneficial effects of cognitive training as a preventative treatment for AD. These novel contributions include: 1) the use of a new generation mouse model of AD; 2) a comparison of the impacts of repeated single versus multi-domain cognitive training; 3) a focus on cognitive training in isolation versus in combination with other strategies (environmental enrichment, exercise, pharmacology, etc); 4) the use of various learning and memory tasks as functional assays for different learning and memory networks; 5) the assessment of a multiple pathologies associated with AD in many of these same brain areas.

### 4.2. New generation knock-in mouse model of AD

More traditional mouse models of AD have overexpressed APP or APP and presinilin1 (PS1) which led to accumulation of unusual fragments generated by α-secretase, such as C-terminal fragment-β (CTF-β). CTF-β is more toxic than Aβ and CTF-β does not accumulate in human AD brains. A recent study estimates that most neuropathological features of these earlier mouse models are due to artifacts related to APP overexpression (Saito et al., 2014) and may explain the lack of translational success of all candidate medications tested in clinical trials. We have been using a second-generation AD model recently developed at the Riken (Saito et al., 2014) which has a modified APP gene that has humanized Aβ sequence with three mutations in APP^NL-G-F^. This mouse model produces robust age-related spread of Aβ aggregates and cognitive problems with endogenous levels of APP. Our lab has recently characterized the APP^NL-G-F^ mouse in several experiments and found that these mice display significant Aβ plaque through regions of NC and HPC and display cognitive impairments at 6 months, but not 3 months of age (Mehla et al., 2019a). The 6-month-old APP^NL-G-F^ mice also showed increased astrocytosis in the hippocampus, neocortex, medial septum/diagonal band as well as other brain areas. Other brain changes in APP^NL-G-F^ mice include cholinergic and norepinephrine dysfunction. Although further research is required, the demonstration of successful use of repeated MT in the APP^NL-G-F^ mice in virtually eliminating both brain pathology and associated cognitive impairments associated with AD suggests that research investigating preventative treatments using this mouse model of AD might be more translatable to humans because of its similarity to human AD pathology.

### 4.3. Effects of different forms of cognitive training on various neural networks implicated in learning and memory functions

In an earlier report, we showed that the App^NL-G-F^ mouse model of AD, starting from approximately 6 months of age, have learning and memory impairments and the nature of these cognitive deficits indicate that several key memory networks have been rendered dysfunctional (Mehla et al., 2019a). For example, the App^NL-G-F^ mice showed severe impairments in the acquisition of the spatial version of the MWT. This spatial navigation task is a sensitive assay of a memory network in which the HPC and retrosplenial cortex play central roles (Sutherland et al., 1988). This AD mouse model also shows impairments on the novel object recognition task (NOR), a task shown to be dependent on the perirhinal cortex which is a terminal region of the ventral stream of visual processing (Mumby and Pinel, 1994; Kealy and Commins, 2011). In addition to likely perirhinal dysfunction in the App^NL-G-F^ mouse brain, a neural network centered on the amygdala implicated in aversive and appetitive classical conditioning processes seems compromised as these AD mice were also impaired in cued and context fear conditioning processes (Kapp et al., 1979; Kim et al., 1993; Antoniadis and McDonald, 2000). In addition to compelling behavioral evidence that these memory networks are dysfunctional in this mouse model of AD we also showed in the same subjects that these networks show many of the hallmark pathologies of human AD including amyloid plaques, microglial activation, and cholinergic dysfunction (Mehla et al., 2019a).

In the present report, we used this new foundation of knowledge about the App^NL-G-F^ mouse model of AD and wanted to assess potential preventative treatment approaches. This study focused on indications from epidemiological and clinical studies that cognitive training is a viable preventative approach for this form of age-related cognitive decline.

### 4.4. Effects of different forms of cognitive training on brain pathology associated with AD

Along with the powerful cognitive effects of MT, we found associated decreases in Aβ pathology in various brain regions such as MSDB complex, HPC, RSC, PRhC & CCA of 12 months old App^NL-G-F^ mice exposed to MT. These brain regions have been implicated in various cognitive functions including learning and memory (McDonald and White, 1993; Mumby and Pinel, 1994; Sutherland et al., 1988; Bird and Burgess, 2008; Buckley, 2005; Czajkowski et al., 2014). The reduction in the amyloid burden in these areas may be responsible for the improved performance on the various assays of learning and memory function.

Glial cell dysfunction observed in the postmortem human AD brain has been reported in various clinical studies (Nagele et al., 2003; Hashioka et al., 2008). In the present study, we also found that MT caused reduced microgliosis in APP^NL-G-F^ mice. However, ST only showed reduction in microgilosis in specific brain regions. These findings from the present study indicate the superiority of MT over ST.

The MSDB complex, a part of the basal forebrain, is mainly responsible for cholinergic inputs to HPC and progressive deterioration and/or dysfunction of cholinergic cells in basal forebrain in aging and neurodegenerative diseases including AD has been reported in previous studies (Whitehouse et al., 1982; Bierer et al., 1995; Gil-Bea et al., 2005; Roman and Kalaria, 2006; Craig et al., 2011). In a previous study, we also reported the loss of cholinergic neurons in MSDB complex of APP^NL-G-F^ mice (Mehla et al., 2019a). In the present study, we found that MT prevented the loss of ChAT positive cells in MSDB complex in 12 months old APP^NL-G-F^ mice. However, ST did not rescue the loss of cholinergic neurons. These findings indicate that repeated and varied cognitive training can preserve cholinergic neurons required for various cognitive functions.

Based on the pattern of results reported in this series of experiments, it is highly likely that repeated MT ameliorate cognitive deficits in APP^NL-G-F^ mice via reducing brain amyloid pathology, microgliosis and preserving cholinergic function.

### 4.5. Relationship of current findings to the existing literature

The present report is supported by epidemiological studies in which the importance of life-long learning as well as mental and physical exercise in aging and neurological disorders has been documented (see a review by Fillit et al., 2002) as well as correlational human studies indicate that extensive mental exercise can provide protection against AD (Katzman, 1993; Snowdon et al., 1996). Here, we can also say that MT showed superiority over ST to ameliorate cognitive deficits and pathology in AD. This conclusion is also supported by previous clinical studies in which it has been reported that MT program could produce a broader effect on improvement of overall cognitive functions compared with single-domain possibly due to cognitive cooperation across different brain processes (Cheng, et al., 2012; Anguera et al., 2013; Hill et al., 2017; Li et al., 2016). In line with the present findings, the beneficial effect of MT on neuropsychological outcomes such as delayed memory, naming, visuospatial ability, executive functions, and attention of patient with early stage of AD has been reported recently in a clinical study (Nousia et al., 2018) which support the use of various cognitive tasks in a repeated manner in the present study. However, repeated ST did not prevent the development of learning and memory deficit in 12 months old App^NL-G-F^ mice on the same task. In contrast to these results, previous studies reported that repeated training in the MWT ameliorates memory deficits in 3xTg-AD mice (Billings et al., 2007; Martinez-Coria et al., 2015). The discrepancies in the results may be due to use of APP-KI mouse (APP^NL-G-F^) model of AD which is different from 3xTg-AD mouse model and experimental design.

Several nonpharmacological strategies for reducing AD pathology and associated cognitive impairments have been evaluated including cognitive training, cognitive stimulation and cognitive rehabilitation/ enrichment, and enhancing neuroplasticity (Woods et al., 2012; Bahar-Fuchs et al., 2013; Herholz et al., 2013; Dardiotis et al., 2018). In an interesting systematic review, it has been suggested that cognitive training is most effective when compared to other nonpharmacological approaches (García-Casal et al., 2017). This view is supported by the present results in which we show clear causal evidence that certain forms of cognitive training, in isolation, eliminates cognitive impairments and brain pathology associated with AD. Specifically, we showed that repeated MT but not ST produced these dramatic effects using a novel rodent model- of AD, the APP^NL-G-F^.

## 5. Summary and Future Directions

The present experiments provide strong causal evidence that multi-domain cognitive training reverses severe cognitive impairments and associated pathological changes in the brain of a new generation knock-in model of AD. These effects indicate that repeated cognitive training that engages multiple learning and memory networks will be these most effective at reversing neurodegenerative processes. It remains to be seen whether these effects are due in fact to learning different things, or to the influence of general enrichment associated with the different tasks. Exposure to different contexts in the absence of cognitive tasks need to be compared to MT and ST groups to answer this question. Importantly though these data do demonstrate that the exercise and enrichment garnered from just repeated MWT training (APP-ST) is not sufficient to rescue memory. Further, the present results combined with our analysis of the existing research literature suggests that implementing MT in combination with other lifestyle factors like voluntary physical exercise or diet could be an effective non-pharmacological approach to prevent or delay the progression of Alzheimer disease. Future work will be directed at assessing the effects of other lifestyle preventative measures alone or in combination with MT.

## Acknowledgements

This work was supported by Natural Sciences and Engineering Research Council of Canada (NSERC) Discovery Grant #40352, #06347, and #03857 to MHM, RJM, and RJS respectively, Alberta Innovates (MHM), Alberta Alzheimer Research Program (MHM), Alzheimer Society of Canada (MHM, RJM), Alberta Prion Research Institute (MHM, RJS), and Canadian Institute for Health Research (MHM, RJM). We thank Dr. Takashi Saito and Prof. Takaomi C Saido from Laboratory for Proteolytic Neuroscience RIKEN Center for Brain Science, Wako-shi, Saitama, Japan” for providing the App^NL-G-F/NL-G-F^ mice as a gift. We also thank Di Shao for animal breeding.

## Author contributions

JM, MHM, RJM designed and conceptualized the experiments. JM, HS, SGL, and SHD performed the behavioural experiments. JM analyzed the behavioural data. JM performed the immunohistochemistry. JM & HK analyzed the immunohistochemistry data. JM, RJM and MHM wrote the manuscript, which all authors commented on and edited. MHM, RJM and RJS provided the resources. MHM and RJM supervised the study.

---

**Supplementary Fig. 1:**
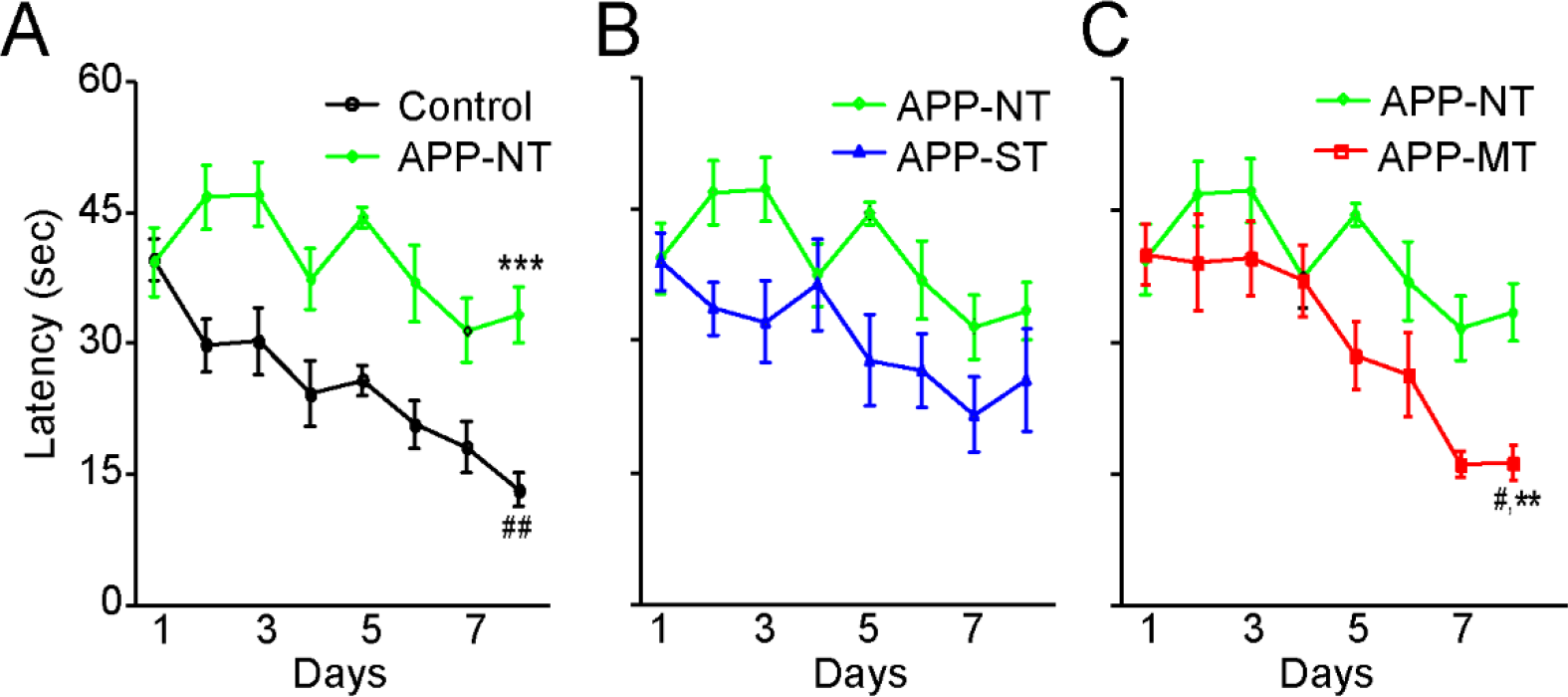
Comparative analysis of escape latency of 12 months old APP^NL-G-F^ mice exposed to different interventions in hidden platform version in MWT. (A) Comparison of escape latency of control and APP-NT groups. (B) Comparison of escape latency of APP-NT and APP-ST groups. (C) Comparison of escape latency of APP-NT and APP-MT groups. Data is presented as mean ± SEM. *,#p < 0.05; **,#p < 0.01; ***p < 0.001; ***- as compared with control group on day 8; ##- as compared with day 1 for control group; #- as compared with day 1 for APP-MT group; **- as compared with APP-NT on day 8. Control group- C57BL/6. APP-NT group- APP^NL-G-F^ with no training. APP-MT group- App^NL-G-F^ mice exposed to multi-domain cognitive training (MT). APP-ST group- App^NL-G-F^ mice exposed to single-domain cognitive training (ST).

**Supplementary Fig. 2:**
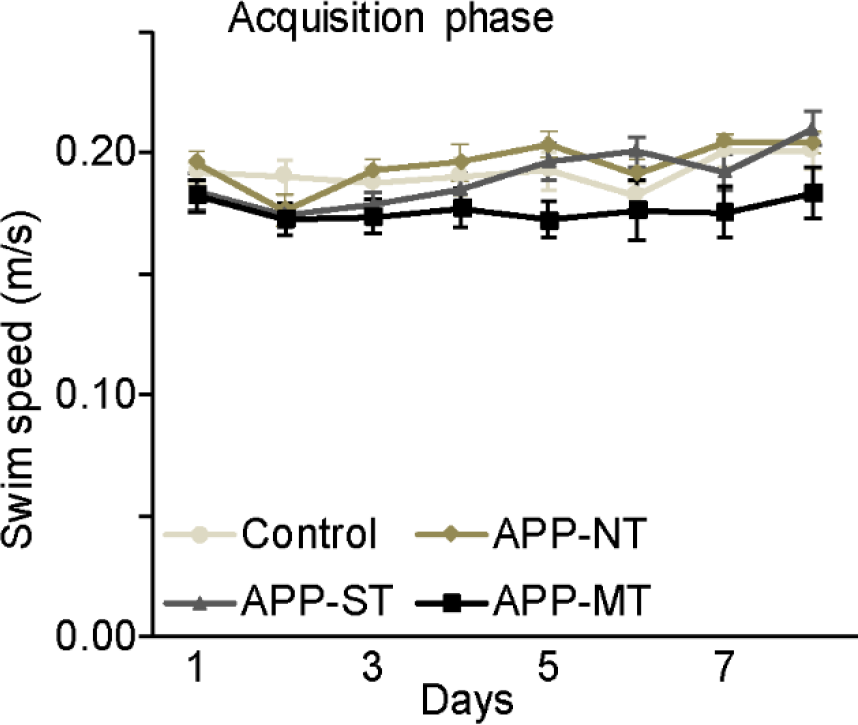
Effects of cognitive training on swim speed of 12 months old mice in the MWT. Data is presented as mean ± SEM. Control group- C57BL/6. APP-NT group- App^NL-G-F^ with no training. APP-MT group- App^NL-G-F^ mice exposed to multi-domain cognitive training (MT). APP-ST group- App^NL-G-F^ mice exposed to single-domain cognitive training (ST).

**Supplementary Fig. 3:**
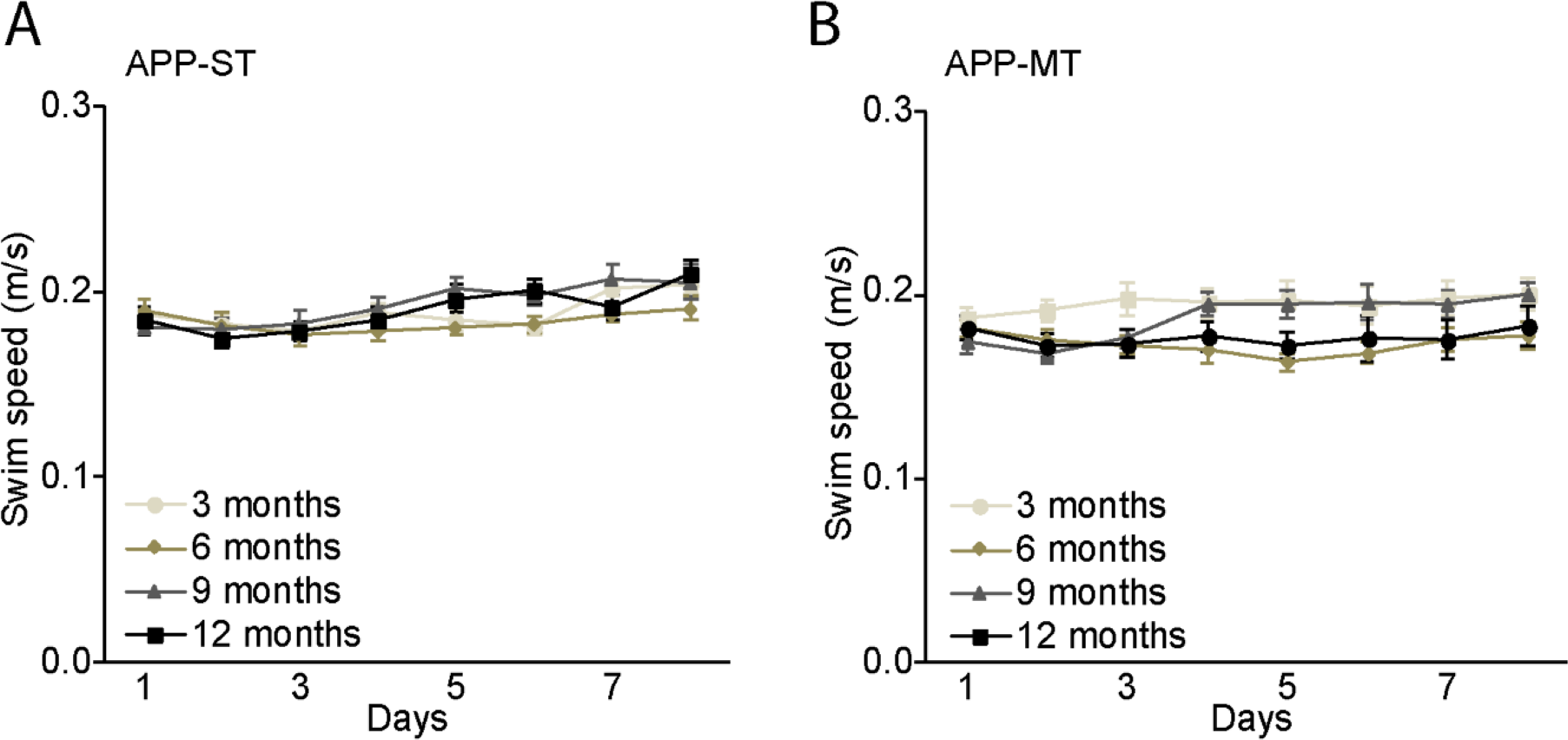
Age dependent swim speed of App^NL-G-F^ mice exposed to single (A) and multi-domain cognitive training (B) in the MWT. Data is presented as mean±SEM. APP-ST group- App^NL-G-F^ mice exposed to single-domain cognitive training. APP-MT group- App^NL-G-F^ mice exposed to multi-domain cognitive training. Data is presented as mean ± SEM.

**Supplementary Fig. 4:**
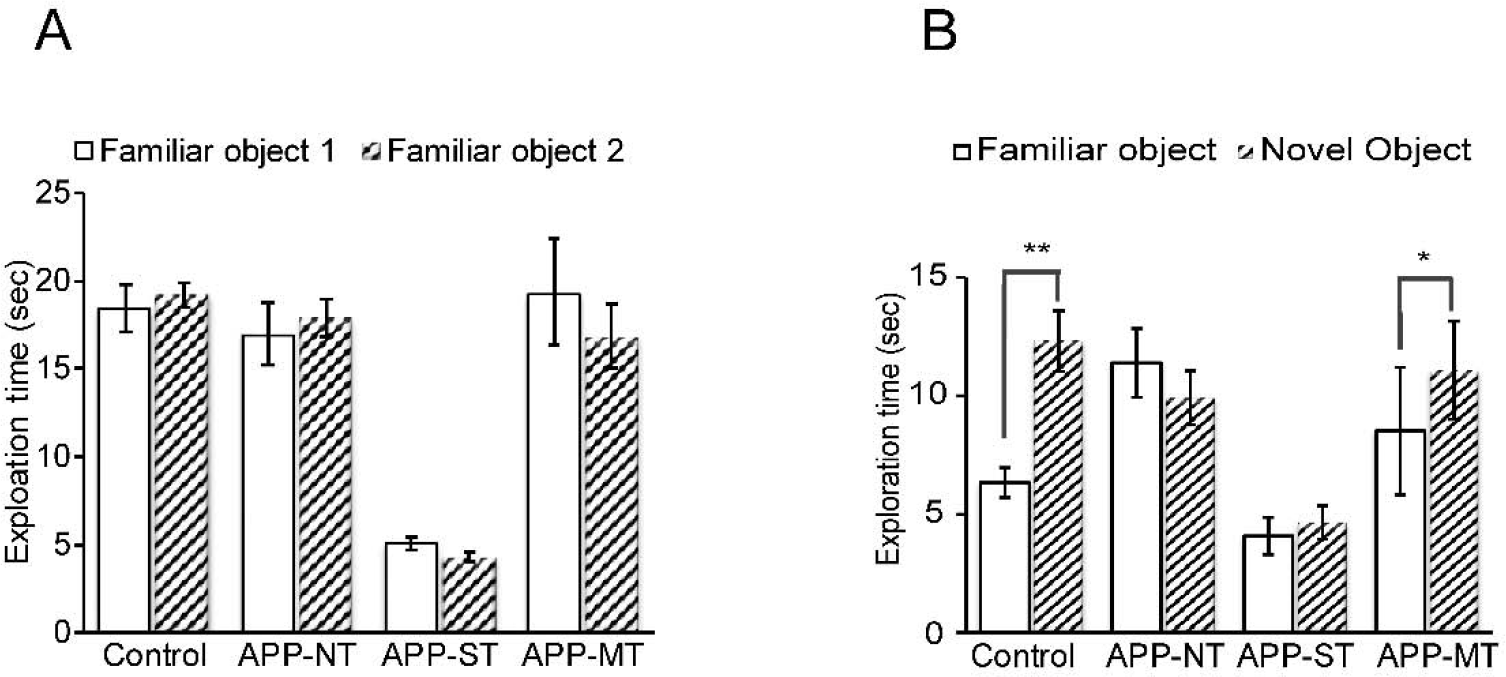
Effects of cognitive training on exploration time of 12 months old mice in object recognition test. Data is presented as mean ± SEM. Control group- C57BL/6. APP-NT group- APP^NL-G-F^ with no training. APP-MT group- APP^NL-G-F^ mice exposed to multi-domain cognitive training (MT). APP-ST group- APP^NL-G-F^ mice exposed to single-domain cognitive training (ST).

